# An fMRI dysmaturity signature in preterm neonates: Responsiveness of brain areas and its relation to newborn brain development

**DOI:** 10.64898/2026.09.24.754103

**Authors:** Martina Attenni, Gianni Valerio Vinci, Maria Luisa Scattoni

## Abstract

Early brain development is intrinsically dynamic, yet most neonatal resting-state functional magnetic resonance imaging (rs-fMRI) studies rely on static and symmetric measures of functional connectivity. Here, we investigated whether response-based metrics can capture latent dynamical organization of the neonatal brain.

We analysed rs-fMRI data from the Developing Human Connectome Project (*n* = 714). Using a statistical-physics framework grounded in the fluctuation–dissipation theorem, we estimated intrinsic pairwise response functions between 46 cortical and subcortical grey-matter regions, enabling quantification of regional responsiveness and response duration.

Response-based measures differed markedly from conventional functional connectivity and predicted gestational age at birth from a single scan. Preterm infants showed reduced responsiveness and shorter response persistence compared with term-born neonates. Importantly, directed analyses revealed a developmental reorganization of driver–listener roles across the brain, with preterm infants preserving the regional ordering of the term hierarchy but expressing it at approximately half amplitude.

These findings reveal that intrinsic responsiveness captures a previously underappreciated dynamical dimension of neonatal brain organization and provides sensitive markers of neurodevelopmental maturity. Response-based metrics may offer a principled framework for detecting early network dysmaturation following preterm birth. They further indicate that prematurity does not reorganize the directed hierarchy of the neonatal connectome so much as arrest its expansion.

## 1 Introduction

Brain development is a complex cascade of adaptive, tightly coordinated processes shaped by genetic programs and continuously modulated by a changing environment. This dynamic process unfolds across multiple developmental stages, each characterized by distinct cellular and structural transformations. During the embryonic and early fetal periods, rapid neurogenesis, neuronal migration, and the initial patterning of major cortical and subcortical structures lay the foundation for large-scale neural architecture [1]. The evolving gross anatomy of the fetal brain corresponds to extensive and dynamic changes at the cellular scale, including differentiation of neural progenitors, establishment of transient developmental layers, and formation of early thalamocortical and corticothalamic pathways. As gestation progresses, the fetal brain undergoes pronounced growth, beginning as a smooth lissencephalic structure and gradually acquiring characteristic gyral and sulcal complexity through orderly, stage-specific folding processes, accompanied by maturation of gray and white matter compartments, exuberant synaptogenesis, and the emergence of early functional circuits [1]. Postnatally, the brain continues to develop through prolonged myelination, synaptic pruning, and refinement of large-scale networks, reaching approximately 90% of adult volume by early childhood while structural and functional maturation extend into adolescence [1, 2]. Together, these developmental cascades illustrate that brain maturation is inherently dynamic, driven by time-dependent interactions between genetic patterning, cellular events, and environmental input. This perspective underscores the need for analytical approaches capable of capturing temporal and functional dynamics rather than relying solely on static measures of brain organization. Over the past decades, the growing emphasis on individualized approaches to diagnosis and treatment has driven the development of increasingly sophisticated investigative methods in human neuroscience.[3][4] Inter-individual variability in cognition and behavior is closely reflected in the structural organization of the brain, with non-invasive MRI capturing meaningful anatomical differences across individuals and linking them to behavioral phenotypes [5]. More recently, advances in network neuroscience have shown that individual differences also emerge from the dynamic reconfiguration of functional brain networks, with dynamic connectivity measures capturing how regions shift between functional communities over time and how these temporal fluctuations relate to cognitive, social, and emotional functioning [6]. Characterizing brain function across early gestational stages provides crucial insight into the fundamental processes that shape later developmental trajectories, while recent evidence indicates that distinct functional connectivity patterns can capture biologically and clinically relevant heterogeneity in neurodevelopmental disorders, including autism[7] [8, 9]. Resting-state functional MRI (rs-fMRI) has enabled the identification of coherent resting-state networks (RSNs) in the fetal and neonatal periods, revealing that primary sensorimotor, auditory, and visual networks are already detectable around term-equivalent age, whereas higher-order association networks remain fragmented and only gradually converge toward adult-like topology in infancy and early childhood [10–13]. Functional connectivity (FC), typically quantified as the temporal correlation of blood oxygen level–dependent (BOLD) activity between regions, has been the predominant tool for studying emerging network architecture in this age range [14, 15]. However, FC is inherently static and symmetric, summarizing covariation patterns without explicitly capturing how activity in one region evolves over time in response to changes in another [16]. Preterm birth, which exposes the immature brain to the extrauterine environment during a period of rapid growth and heightened plasticity, is associated with increased risk for long-term cognitive, motor, and psychiatric difficulties even in the absence of overt structural brain injury [17–19]. Within the Developing Human Connectome Project (dHCP), high-resolution rs-fMRI studies have shown that preterm infants scanned at term-equivalent age exhibit widespread reductions in FC across multiple RSNs compared with term-born peers, alongside focal increases in parieto-motor connectivity that may relate to later motor and visuospatial difficulties [20]. These findings indicate that prematurity alters the maturation of large-scale functional networks in a dose-dependent manner and that perinatal FC patterns carry information about subsequent neurodevelopmental vulnerability [21]. Considering brain development as a highly dynamic process, metrics that capture temporal evolution are essential for understanding maturation and for distinguishing typical from altered trajectories. In this context, responsiveness provides a complementary framework to conventional FC, as it explicitly quantifies how neural activity in one region would change over time in response to perturbations in another, thereby incorporating both directionality and temporal structure [22, 23]. By estimating intrinsic responsiveness directly from rs-fMRI time series, it becomes possible to probe latent dynamical properties of neonatal brain networks that static connectivity measures may overlook. This approach aligns closely with the dynamic nature of early brain development, emphasizing the central role of temporal trajectories in shaping emerging functional organization and neurodevelopmental outcomes. The Developing Human Connectome Project (dHCP), funded by the European Research Council, provides an open-access resource of structural and functional MRI acquired between 20 and 44 weeks postmenstrual age using a dedicated neonatal imaging platform [24–26]. This large-scale dataset enables 4D connectivity mapping to track perinatal brain developmental trajectories under both typical and adverse conditions, improving understanding of normal maturation and supporting earlier detection and intervention for neurological and psychological disorders [20]. Building on this framework, the present study applies a response-based statistical-physics approach to rs-fMRI data from the dHCP to quantify regional responsiveness and response duration in newborns from preterm to term age, and to determine how these dynamical markers relate to gestational age at birth, postmenstrual age at scan, and conventional FC.

## 2 Materials and Methods

### 2.1 Participants

We analysed 714 neonatal rs-fMRI datasets from the third data release of the Developing Human Connectome Project (dHCP, release 3), including both term-born and preterm-born infants [24]. Term infants (n = 519) were born at a median gestational age (GA) of 40.1 weeks (IQR 39.0–40.9) and were scanned soon after birth at a median postmenstrual age (PMA) of 41.3 weeks (IQR 40.1–42.6). Preterm infants (GA at birth *<* 37 weeks; n = 195) were born at a median GA of 32.6 weeks (IQR 29.4–35.1) and scanned at a median PMA of 36.9 weeks (IQR 35.0–40.1); of these, 32 were extremely preterm (*<* 28 weeks), 55 very preterm (28–32 weeks) and 108 moderate-to-late preterm (32–37 weeks). The final sample comprised 385 males and 329 females. Where a single scan per infant was required, the acquisition closest to 38 weeks PMA was retained; 89 infants contributed two sessions and were used for the longitudinal analyses. Inclusion and exclusion criteria, as well as details of successive exclusion steps, are reported in the dHCP neonatal data release documentation [24]. All infants were recruited at St Thomas’ Hospital, London, and imaged at the Evelina Newborn Imaging Centre with written informed parental consent, under approval of the UK National Research Ethics Authority (14/LO/1169).

### 2.2 fMRI acquisition

Resting-state fMRI data were acquired as part of the dHCP protocol on a 3 T Philips Achieva system equipped with a custom 32-channel neonatal head coil and a dedicated neonatal imaging set-up optimised for unsedated scanning [25, 26]. Infants were scanned during natural sleep, with standardized positioning, multi-layer hearing protection and continuous monitoring of heart rate, oxygen saturation and temperature throughout the examination [24, 26]. Blood-oxygen-level-dependent (BOLD) fMRI was collected using a multiband 9× accelerated gradient-echo EPI sequence (TR = 392 ms, TE = 38 ms, flip angle 34^◦^, voxel size 2.15 2.15 2.15 mm^3^, 45 slices, 2,300 volumes, total acquisition time ≈ 15 min), as described in the dHCP functional protocol [24, 25]. High-resolution T2-weighted and T1-weighted anatomical images were also acquired for morphometric analysis and registration [24].

### 2.3 Data processing

All rs-fMRI datasets were preprocessed using the dedicated dHCP neonatal pipeline [24, 25]. Briefly, preprocessing comprised correction of susceptibility-induced distortions using reversed-phase-encoding spin-echo images, slice-to-volume motion correction and dynamic distortion correction, rigid-body realignment, and regression of residual motion-, multiband- and cardiorespiratory-related artefacts [25]. Datasets with excessive motion (more than 10% volumes flagged as motion outliers based on framewise displacement) or clinically significant incidental MRI findings, according to the radiology scoring in the dHCP database, were excluded [24]. T2-weighted images were segmented into tissue classes and non-linearly registered to the dHCP neonatal template to propagate atlas labels to native space [24, 27]. From the minimally preprocessed rs-fMRI data, we extracted BOLD time series for 87 cortical and subcortical regions of interest (ROIs) defined in the dHCP neonatal parcellation, averaging voxelwise signals within each ROI in grey and deep grey matter [24].

### 2.4 Response-based metric

The Fluctuation-Dissipation Theorem (FDT) is a standard result from statistical physics [28]. In its most general formulation it allows one to derive the time dependent response *R_ij_*(*t*) of one variable *i* to an instantaneous kick of the variable *j* performed at initial time *t* = 0, from non-linearly weighted correlation functions of the unperturbed system. This result is particularly relevant whenever a perturbation experiment is not feasible, as in the case of human brain imaging, but sufficient observations of the unperturbed system are available which in our case is the resting-state fMRI signal. For general non-linear system, assuming stationarity and Markovianity, the theorem predicts *R_ij_*(*t*) = *x_i_*(*t*)*∂_x_ p*(*x*_0_) where *p*(*x*_0_) is the initial distribution which coincides with the stationary measure and the average is done over unperturbed trajectories. While brain dynamics are in general non-linear, the fMRI signal of a region of interest (ROI) results from averaging the activity of many spatially neighbouring neurons, so that by the central limit theorem the resulting signal is expected to be approximately Gaussian. There is also ample evidence that the resting state in particular is well described by linearised dynamics around a stable fixed point [29, 30].

When the system is linear the FDT greatly simplifies [31, 32]. For a zero-mean Gaussian stationary state with covariance *C* =(*xx*^T^) the measure is 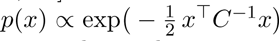 so that 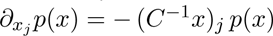 Substituting into the general expression above, the response function collapses onto a purely second-order object,

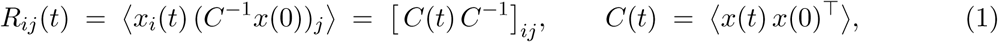

where *C*(*t*) is the lagged covariance matrix, notably *C*(0) = *C* is the functional connectivity matrix typically reported in the literature. Equation (1) is the operational core of this work: the response of the system to a perturbation that was never applied is recovered entirely from the covariance and the lagged covariance of the spontaneous signal. Unlike *C*, the matrix *R* is in general *not* symmetric, and *R_ij_* = *R_ji_* carries the directional information that functional connectivity discards.

#### Estimation from resting-state data

For each subject the regional BOLD signal was arranged in a matrix *X* ∈ R*^n^*^×*T*^ and each row was independently z-scored, *X_i,t_* ←(*X_i,t −_μ_i_*)/(*σ_i_* + *ɛ*) with *ε* = 10^−6^, so that *C* has unit diagonal and all regions contribute on a common scale. The instantaneous and one-step lagged covariances were estimated as

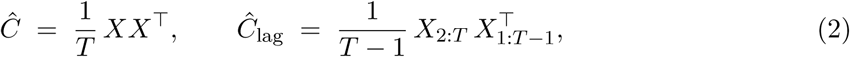

and the response matrix at one repetition time followed from Eq. (1),

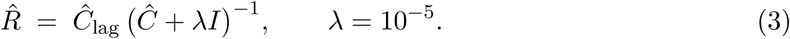

The Tikhonov term *λI* stabilises the inversion against the near-singular directions of *C*^^^ that arise when regions are strongly collinear. The entry *R*^^^*_ij_* is read as the response of region *i*, one TR later, to an instantaneous unit perturbation of region *j*, after the instantaneous covariance structure has been projected out.

Of the 87 parcels of the neonatal atlas, all analyses were restricted to the 46 cortical and subcortical grey-matter regions (23 left, 23 right) that remain after excluding white matter, CSF, ventricles, corpus callosum, brainstem, cerebellum and background. Regions were reordered into left- and right-hemispheric blocks, and self-connections were removed by setting the diagonal of *R* to zero before computing any node-level quantity.

#### Continuous-time dynamics and response duration

Because *R*^^^ describes propagation over one sampling interval, it relates to the underlying continuous-time dynamics through *R*(Δ*t*) ≈ exp(*J* Δ*t*), with *J* the Jacobian of the linearised system. Expanding to first order gives the estimator

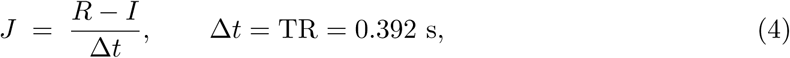

which is accurate when the eigenvalues of *R* lie close to unity, i.e. when the dynamics evolve slowly relative to the sampling rate. This condition holds for the slow modes analysed here – the leading eigenvalue of *R* has median 0.87 across subjects, and exactly the five modes we track exceed 0.5 – but not for the fast bulk of the spectrum, whose eigenvalues cluster near zero. Appendix C compares the resulting timescales against the exact matrix-logarithm solution. The eigenvalues *λ_k_* of *J* are generally complex; Re(*λ_k_*) *<* 0 identifies a stable, decaying mode, and its magnitude sets the intrinsic timescale over which a perturbation along that mode survives,

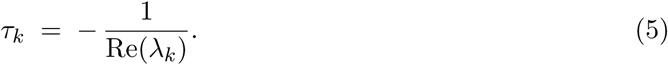

Eigenvalues approaching zero from below therefore correspond to slowly decaying, long-lived modes: what we refer to as increased *response duration*. The *K* = 5 modes with the largest (least negative) real part were tracked per subject and summarised within each age bin.

#### Node-level response metrics

Three scalar summaries were derived per region from the diagonal-free *R*. Recalling that column *j* collects the influence emitted by region *j* and row *i* the influence it receives,

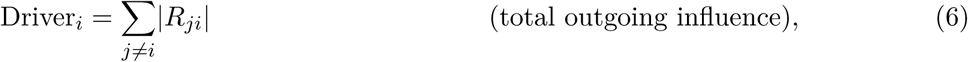

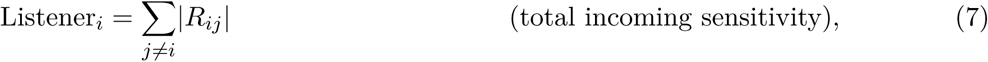

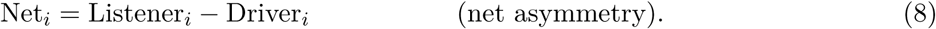

A negative Net*_i_* marks a region that emits more directed influence than it absorbs – a broadcaster or *driver* – whereas a positive value identifies a net *listener*. This driver/listener decomposition has no counterpart in symmetric functional connectivity, where the two sums coincide by construction.

Hemispheric specialisation was quantified, separately for incoming and outgoing strength, by the asymmetry index of each of the 23 bilateral region pairs,

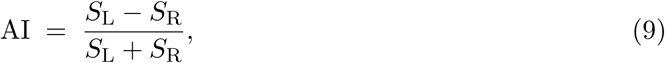

with *S* the corresponding directed strength, so that negative values denote left dominance and positive values right dominance.

#### Matrix-level metrics

Two further quantities summarise whole-matrix organisation. The rate at which network structure is remodelled between consecutive age bins was measured by the relative Frobenius-norm change of the group-mean matrix *M̄_t_* ∈ {*C̄_t_, R̄_t_*},

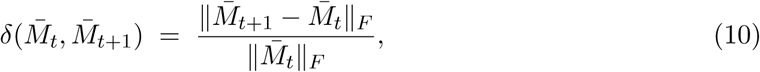

and inter-individual heterogeneity within a bin by the Frobenius distance of each subject’s matrix from the bin centroid, *d_i_* = ∥*M_i_*− *M̄* ∥*_F_*.

#### Response-derived feature set

For the prediction analyses each subject was represented by 161 features computed from a single scan: the 46 outgoing and 46 incoming strengths; 30 principal components of the 1035 unique off-diagonal entries of *R*; the 10 leading eigenvalue magnitudes of *R*; the 23 left–right differences in net asymmetry; three graph-theoretic scalars (global efficiency, weighted clustering coefficient and the Fiedler value of the normalised Laplacian of the symmetrised *R*); and postmenstrual age at scan, sex and the motion metric as covariates.

### 2.5 Statistical analysis

All analyses were carried out in Python (NumPy, SciPy, scikit-learn, statsmodels). Unless stated otherwise, two-sided tests were used and the significance threshold was set at *α* = 0.05.

#### Age binning and group summaries (**Fig. 1**)

Scans were assigned to nine consecutive postmenstrual age bins (≤ 30, 30–32, …, 44–46 weeks). Eigenvalues of *J* and explained variances of *C* were computed per subject and summarised within each bin as mean ± 1.96 SE, an interval that assumes approximate normality of the across-subject distribution and is therefore best supported in the better-populated bins (*n* ≥ 30). The relative Frobenius change between adjacent bins was accompanied by a non-parametric bootstrap: subjects were resampled with replacement independently within each of the two bins, the group means recomputed and the statistic reevaluated over *B* = 1000 resamples, with the 2.5th and 97.5th percentiles giving asymmetric 95% confidence intervals. This procedure makes no distributional assumption and tolerates the strongly unequal bin sizes.

**Figure 1:**
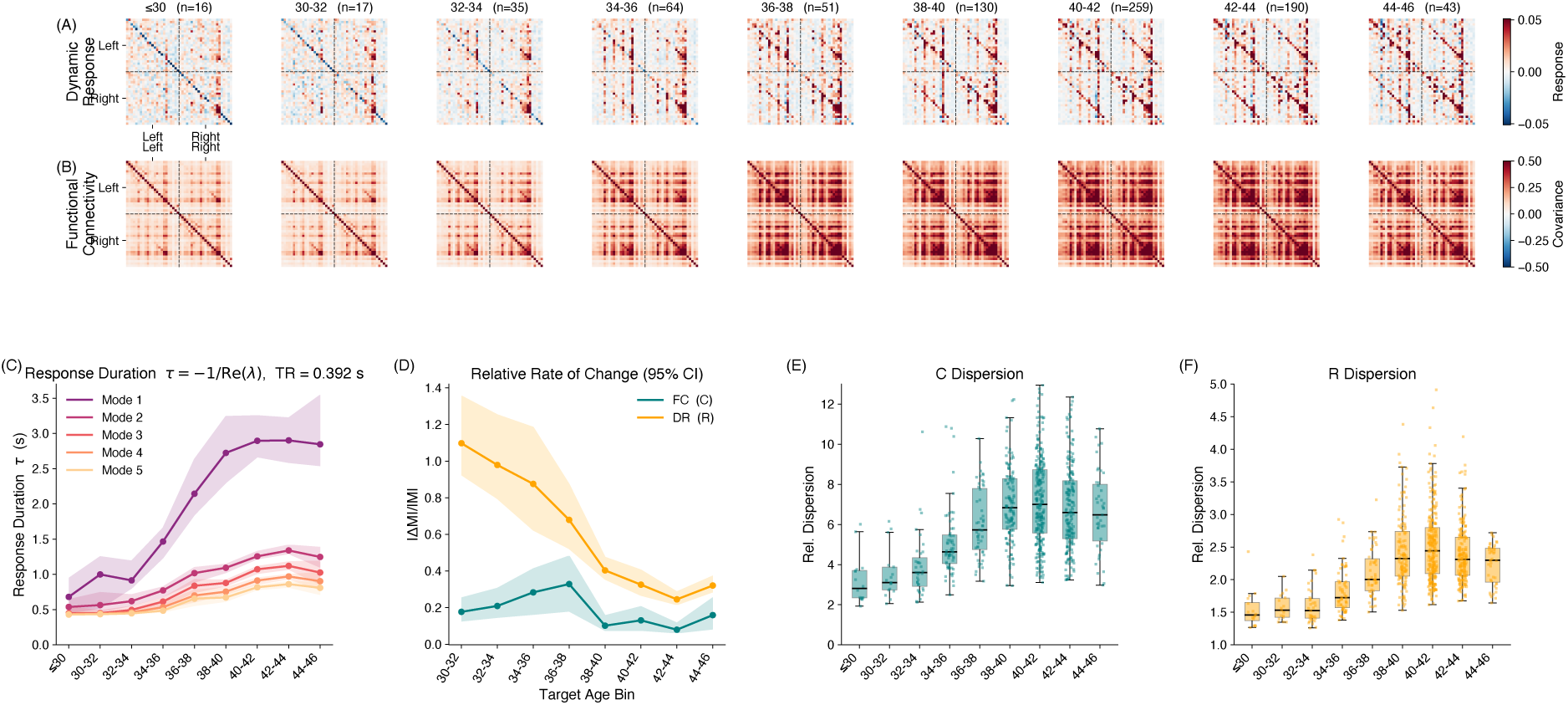
Developmental evolution of functional connectivity and intrinsic response across the neonatal period. **(A, B)** Group-averaged response matrices *R* (A) and functional connectivity matrices *C* (B) in nine postmenstrual age bins from ≤ 30 to 44–46 weeks; the number of scans contributing to each bin is given above the corresponding column. The 46 grey-matter regions are ordered into left- and right-hemispheric blocks, separated by dashed lines. Colour encodes magnitude (red positive, blue negative), with the response scale clipped at the 97th percentile of *R* and the covariance scale fixed at ±0.5. Whereas *C* saturates towards a densely correlated pattern, *R* retains a sparse, markedly asymmetric structure that becomes progressively more organised with age. **(C)** Intrinsic response duration *τ* = −1*/*Re(*λ*) of the five leading modes of the linearised Jacobian *J* = (*R I*)*/*Δ*t*, Δ*t* = TR = 0.392 s, shown as the median across subjects with a bootstrap 95% confidence interval (*B* = 2000). The median rather than the mean is reported because *τ* is a non-linear transform of the eigenvalue and a minority of infants have a non-decaying leading mode (Re(*λ*) ≥ 0, at most 18 of 259 in any bin), for which *τ* diverges. The slowest mode lengthens from 0.68 to 2.85 s between ≤ 30 and 44–46 weeks, a 4.2-fold increase, and all five modes lengthen monotonically with postmenstrual age. **(D)** Relative rate of change of the group-mean matrix between consecutive bins, ∥*M̄_t_*_+1_− *M̄_t_*∥*_F_* / ∥*M̄_t_*∥*_F_*, for *C* (teal) and *R* (orange); bands are 95% bootstrap confidence intervals (*B* = 1000). The response matrix is remodelled several times faster than functional connectivity in the early preterm period, and the two converge by term. **(E, F)** Inter-subject dispersion within each bin, measured as the Frobenius distance of each subject’s matrix from the bin centroid, for *C* (E) and *R* (F). Boxes show median and interquartile range; dots are individual scans.

#### Directed flow and longitudinal change (**Fig. 2**)

Standard errors on driver, listener and net strength were obtained by computing each scalar per subject first and then dividing the across-subject standard deviation by 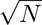, rather than by propagating entry-wise variances, which would assume independence between matrix entries. Longitudinal change was assessed in the 89 infants scanned twice, by testing the within-subject difference (S2 S1) against zero with the Wilcoxon signed-rank test; a one-sample *t*-test was used as a fallback for fewer than ten pairs. These region-wise tests were not corrected for multiple comparisons and are reported as exploratory.

**Figure 2:**
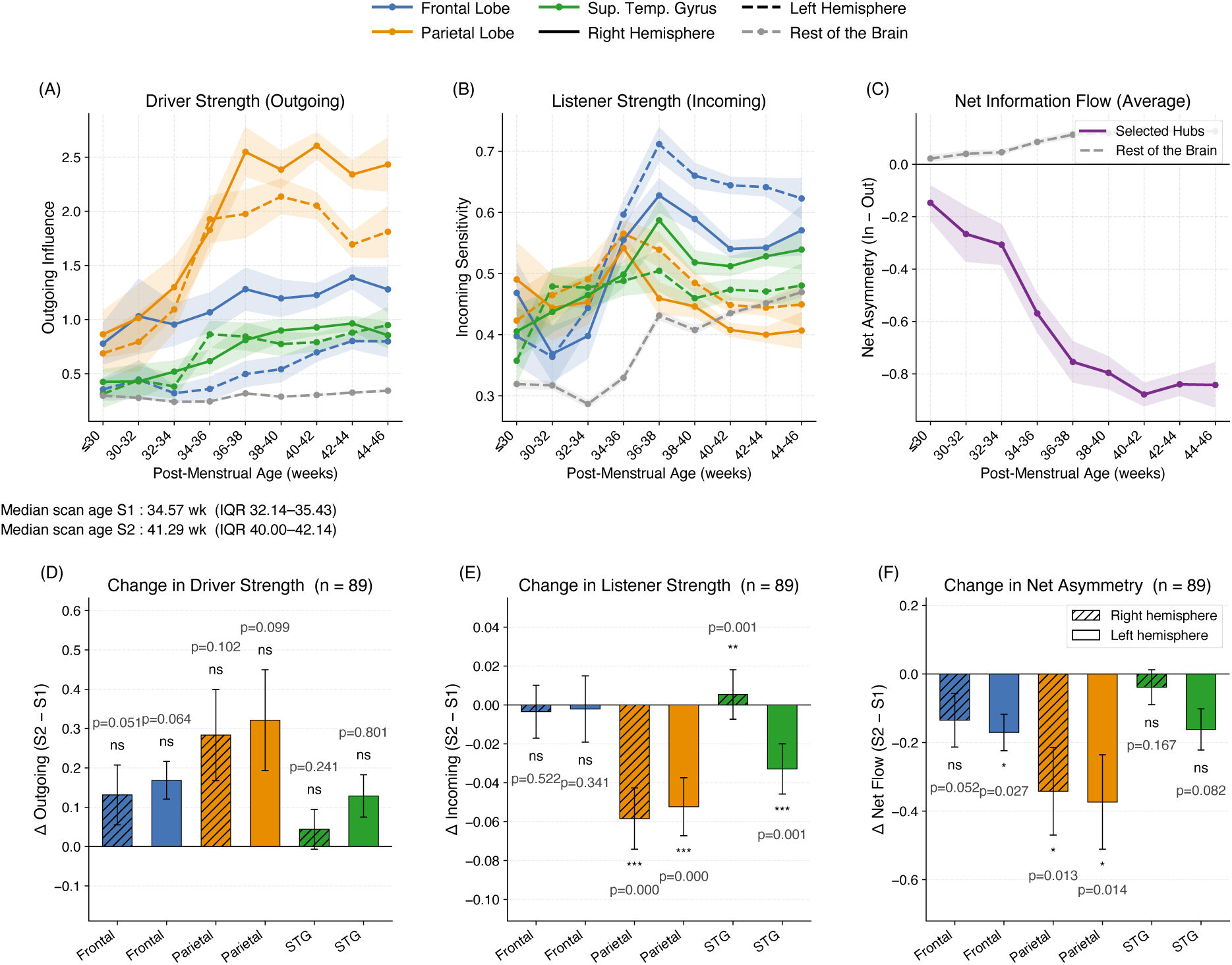
Emergence of directed driver–listener organisation and its longitudinal maturation. **(A–C)** Cross-sectional trajectories across the nine postmenstrual age bins for six *a priori* cortical hubs (frontal lobe, parietal lobe and middle superior temporal gyrus, bilaterally) and for the average of all remaining grey-matter regions. Colour denotes anatomy and line style hemisphere; shaded bands are ±1 SEM computed across subjects. **(A)** Driver strength, the total outgoing influence Σ*_i_*|*R_ij_|*.**(B)** Listener strength, the total incoming sensitivity Σ*_i_*|*R_ij_|*. **(C)** Net asymmetry (incoming outgoing), averaged over the six hubs and over the rest of the brain. The hubs diverge steadily towards negative values while the rest of the brain remains weakly positive, indicating that cortical hubs progressively assume the role of broadcasters. **(D–F)** Within-subject change between the first and second scan in the 89 infants imaged twice (median interval 7.3 weeks). Bars show the mean change (S2 S1) with ±1 SEM; hatching marks right-hemispheric regions. Significance of the change from zero was assessed with the Wilcoxon signed-rank test (*^∗^p <* 0.05, *^∗∗^p <* 0.01, *^∗∗∗^p <* 0.001, ns = not significant).

#### Group contrasts at term-equivalent age (**Fig. 3**)

Preterm and term infants were compared on regional net asymmetry using the Mann–Whitney *U* test, avoiding any normality assumption on a bounded, skewed statistic. The resulting *p*-values were corrected across the 46 regions with the Benjamini–Hochberg false discovery rate procedure, and regions with *q <* 0.05 were retained. Effect sizes are reported as Hedges’ *g*, i.e. Cohen’s *d* with the small-sample correction that matters here given the markedly unequal group sizes (*n* = 519 vs. *n* = 124). Confidence intervals on the between-group difference in means were obtained by bootstrap resampling of both groups (*B* = 2000, percentile method).

**Figure 3:**
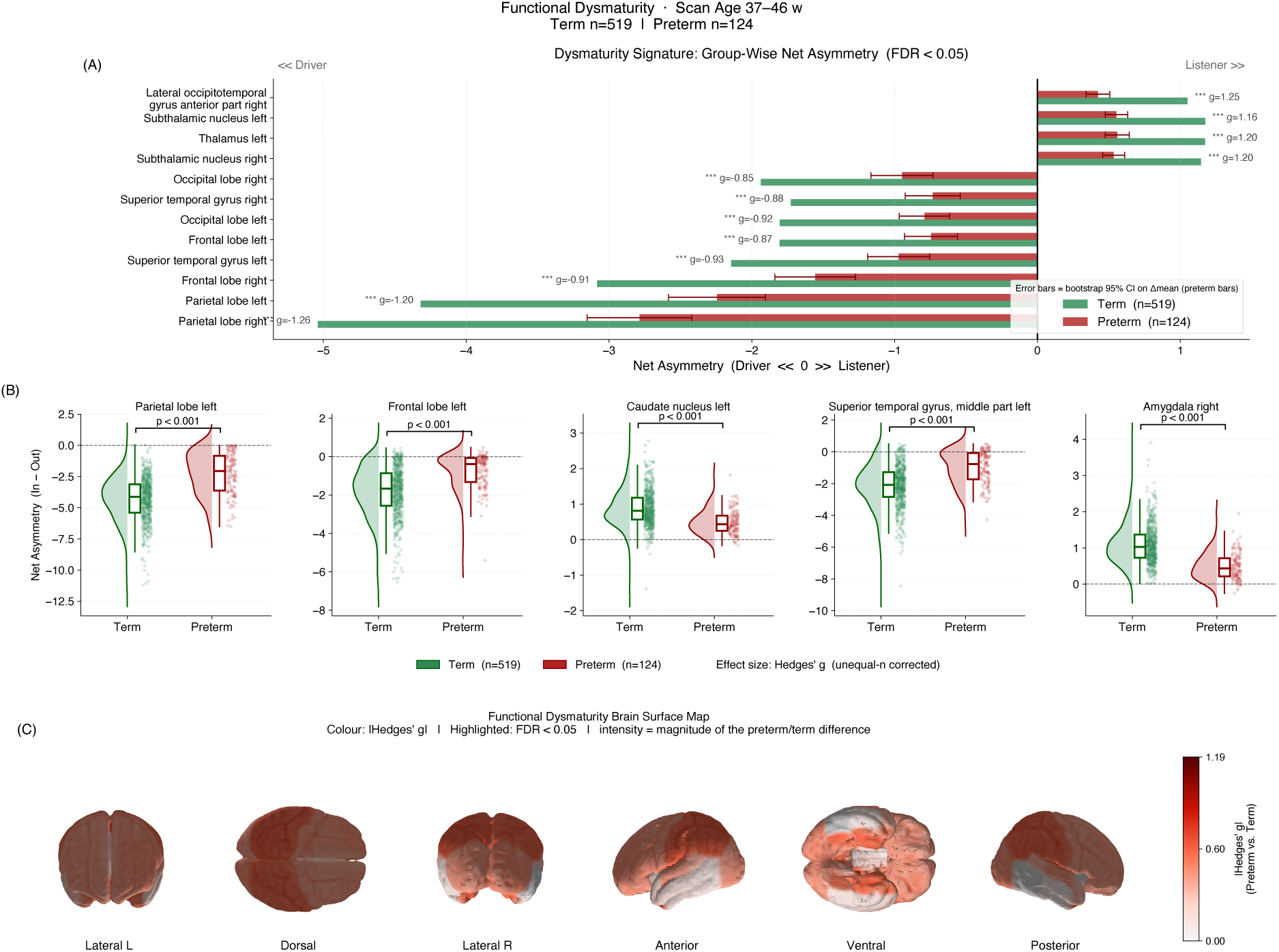
Functional dysmaturity of the response architecture in preterm infants at term- equivalent age. All comparisons use one scan per infant acquired between 37 and 46 weeks postmenstrual age (term *n* = 519, preterm *n* = 124). **(A)** Group-mean net asymmetry for the twelve most strongly differentiating grey-matter regions, of the 44 of 46 surviving Benjamini–Hochberg correction at *q <* 0.05. Negative values denote a driver role and positive values a listener role. Error bars on the preterm bars are bootstrap 95% confidence intervals on the between-group difference (*B* = 2000); Hedges’ *g* is annotated per region. Bilateral homologues appear as pairs throughout. The dominant pattern is a systematic shortfall in the driver role of parietal, frontal, occipital and superior temporal cortex in preterm infants, accompanied by a relative shift towards driving in subcortical and limbic structures. **(B)** Raincloud plots for five representative regions, overlaying a kernel density estimate, a box-and-whisker summary and individual observations; brackets give Benjamini–Hochberg-corrected *p*-values. **(C)** Cortical surface rendering of the effect magnitude, Hedges’ *g*, restricted to regions significant at FDR *<* 0.05; grey denotes non-significant grey matter. Colour encodes magnitude only and is deliberately direction-independent.

#### Lateralisation (**Fig. 4**)

For each bilateral pair, population-level asymmetry was tested against zero with a one-sample *t*-test on the pooled sample, and the birth-timing effect with Welch’s *t*-test, which does not assume equal variances between the term and preterm groups. Because four metrics (incoming and outgoing asymmetry, population and birth effect) were evaluated for every pair, the Benjamini–Hochberg correction was applied jointly across all 4 × 23 tests rather than within each panel, controlling the false discovery rate over the whole display.

**Figure 4:**
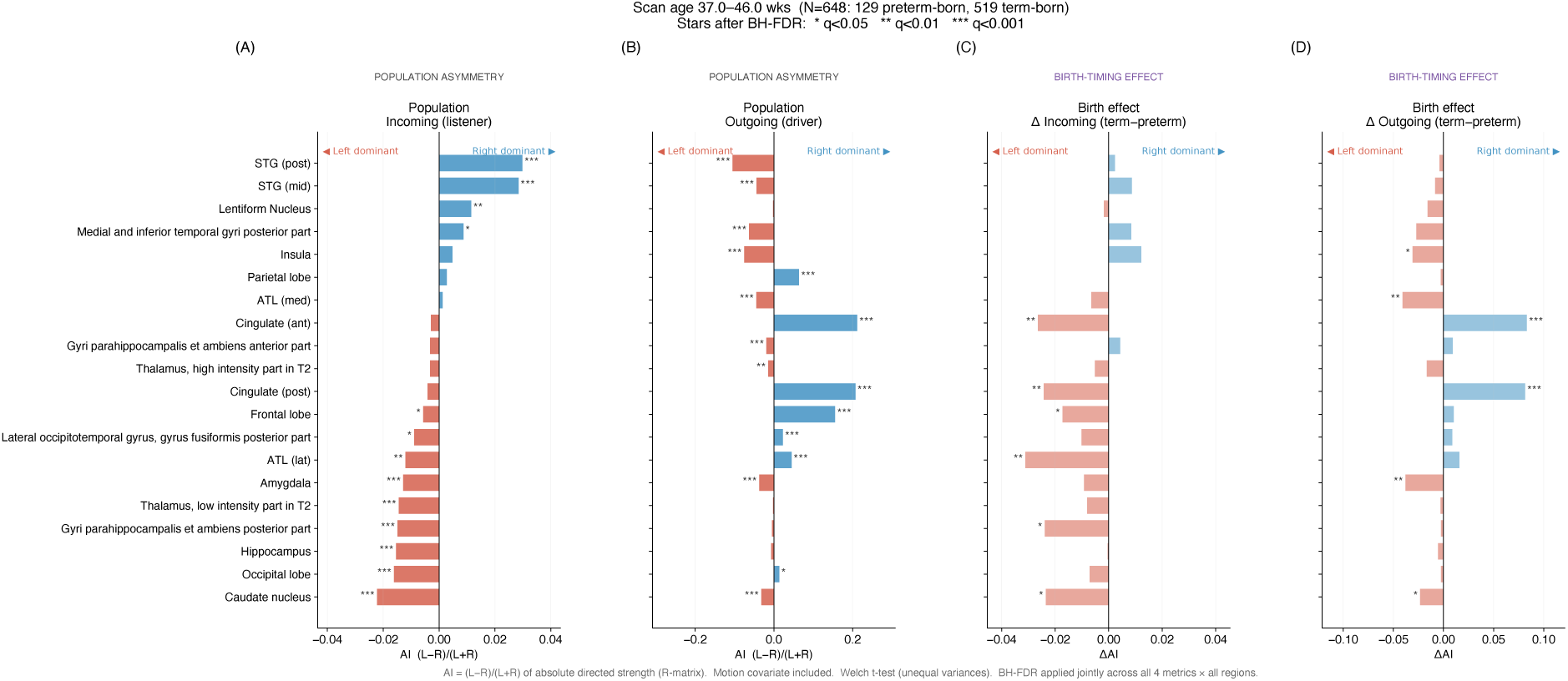
Hemispheric asymmetry of directed strength and its modulation by birth timing. Asymmetry index AI = (*L R*)*/*(*L* + *R*) of absolute directed strength, computed for the 23 bilateral grey-matter pairs in infants scanned between 37 and 46 weeks postmenstrual age (*N* = 648: 129 preterm-born, 519 term-born); the 20 pairs surviving joint Benjamini–Hochberg correction are shown, ordered by incoming asymmetry. **(A, B)** Population-level asymmetry of incoming (listener) and outgoing (driver) strength, tested against zero with a one-sample *t*-test. Red bars denote left dominance, blue bars right dominance. Incoming and outgoing asymmetries frequently point in opposite directions within the same region, a dissociation that is invisible to symmetric connectivity measures. **(C, D)** Birth-timing effect, expressed as the difference in asymmetry index between term- and preterm-born infants (Δ = term preterm) for incoming and outgoing strength, tested with Welch’s *t*-test. Asterisks denote FDR-corrected significance (*^∗^q <* 0.05, *^∗∗^q <* 0.01, *^∗∗∗^q <* 0.001).

#### Prediction of gestational age at birth (**Fig. 5**)

Models were evaluated by 5-fold stratified group cross-validation: folds were stratified on binned birth gestational age so that each fold spans the full range of prematurity, and grouped by subject so that no infant contributed to both training and test sets. Performance is reported as the coefficient of determination and root-mean-square error computed on the pooled out-of-fold predictions, which is a stricter summary than averaging per-fold scores. Feature importance was assessed by permutation on the fitted model (20 repeats), a model-agnostic measure that, unlike impurity-based importance, does not favour high-cardinality features. To separate the contribution of brain organisation from the scan-timing confound, the entire pipeline was rerun with postmenstrual age at scan removed from the feature set; the difference in explained variance between the two models quantifies the unique contribution of scan timing.

**Figure 5:**
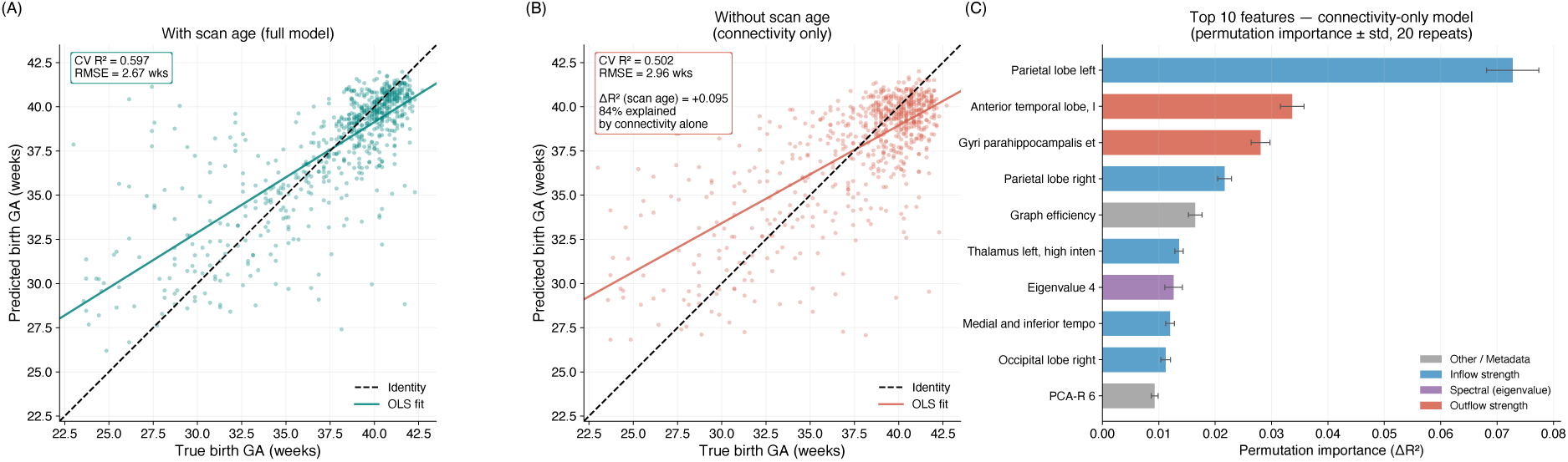
Response-derived features predict gestational age at birth independently of scan timing. **(A)** Out-of-fold predictions of gestational age at birth from the full feature set (*n* = 714, one scan per infant), obtained by 5-fold stratified group cross-validation. The dashed line is identity and the solid line the ordinary least-squares fit. **(B)** The same pipeline with postmenstrual age at scan removed from the feature set. Performance falls only modestly (*R*^2^ = 0.502 vs. 0.597; Δ*R*^2^ = 0.095), so that roughly 84% of the explained variance is carried by the response architecture itself rather than by the timing of the scan. The residual regression towards the mean at low gestational ages is characteristic of the strongly skewed birth-age distribution. **(C)** The ten most informative features of the connectivity-only model, ranked by permutation importance (± standard deviation over 20 repeats) and coloured by feature family. Directed strengths dominate, with a spectral feature entering the top ten, indicating that both the regional balance of outgoing and incoming influence and the global timescale structure of the response carry information about prematurity.

## 3 Results

### 3.1 Developmental evolution of intrinsic response

Group-averaged response and functional connectivity matrices were computed in nine postmenstrual age bins from ≤ 30 to 44–46 weeks (Fig. 1A,B). Both matrices become progressively more structured with age, but they do so at very different rates. Between the two earliest bins the group-mean response matrix changed by 110% of its own norm against 18% for functional connectivity, a factor of 6.2; the gap narrows to roughly twofold by term-equivalent age (Fig. 1D).

The intrinsic timescales of the system lengthen over the same interval. The response duration of the slowest mode, *τ* = −1*/*Re(*λ*), rises from a median of 0.68 s at ≤ 30 weeks to 2.85 s at 44–46 weeks, a 4.2-fold increase, and all five leading modes lengthen monotonically, the fifth from 0.43 to 0.81 s (Fig. 1C). Inter-individual dispersion also increased with age, 2.3-fold for *C* and 1.6-fold for *R* (Fig. 1E,F).

### 3.2 Emergence of a driver–listener hierarchy

Outgoing and incoming influence followed dissociable trajectories (Fig. 2A,B). Right parietal driver strength rose from 0.86 to 2.43 across the age range, a 2.8-fold increase, while the average over all remaining grey-matter regions was essentially flat (0.30 to 0.34). Incoming sensitivity instead peaked at 36–38 weeks and then declined, by 25% in right parietal and 12% in left frontal cortex. Net asymmetry of the six hub regions moved from 0.15 to 0.84 while the rest of the brain drifted only from +0.02 to +0.13 (Fig. 2C).

Among the 89 infants scanned twice (median interval 7.3 weeks), no hub showed a significant change in driver strength (all *p* ≥ 0.051), whereas listener strength fell significantly in parietal cortex bilaterally (*p <* 0.001) and changed significantly in both superior temporal gyri (*p* = 0.001); net asymmetry became significantly more negative in parietal cortex bilaterally and left frontal lobe (Fig. 2D–F).

### 3.3 Dysmaturity of the directed hierarchy after preterm birth

At term-equivalent age, 44 of 46 regions differed significantly between term and preterm infants (Mann–Whitney, FDR *q <* 0.05; Fig. 3). In term-born neonates the hierarchy is sharply differentiated: parietal cortex is the strongest driver (net asymmetry 5.04 right, 4.32 left), followed by frontal, occipital and superior temporal cortex, while every subcortical and limbic structure is listener-dominant, including subthalamic nucleus (+1.17), thalamus (+1.17), amygdala (+1.10) and caudate (+0.88). Preterm infants reproduced this entire ordering at approximately half amplitude: parietal cortex reached only 2.79 and the amygdala +0.49. Regressing the preterm on the term group mean across all 46 regions gave a slope of 0.497 with an intercept indistinguishable from zero and *r* = 0.99 (*R*^2^ = 0.98). No region reversed the sign of its net asymmetry, and the only two regions more extreme in preterm infants were non-significant (*q* = 0.96 and 0.63, |*g*| *<* 0.08).

### 3.4 Hemispheric specialisation of directed influence

Twenty of 23 bilateral pairs showed significant lateralisation of directed strength after joint FDR correction (Fig. 4). Seven regions carried incoming and outgoing asymmetries of *opposite* sign, both surviving correction: the posterior superior temporal gyrus listens right-dominantly (AI = +0.030) while driving left-dominantly ( −0.106), and the frontal lobe shows the converse ( −0.006 incoming, +0.156 outgoing).

The anterior and posterior cingulate carried both the largest population-level outgoing asymmetry (AI = +0.213 and +0.208) and the largest birth-timing effect (ΔAI_out_ = +0.083 and +0.082, *q <* 0.0001), with an oppositely signed shift in incoming asymmetry ( 0.027 and 0.024, *q <* 0.01).

### 3.5 Prediction of gestational age at birth

Directed network features predicted gestational age at birth from a single scan with *R*^2^ = 0.597 (RMSE 2.67 weeks) under 5-fold stratified group cross-validation. Removing postmenstrual age at scan from the feature set reduced this only to *R*^2^ = 0.502 (RMSE 2.96 weeks), so 84% of the explained variance is carried by brain organisation rather than scan timing (Fig. 5A,B). The most informative predictors were incoming strength of the parietal lobe bilaterally and outgoing strength of anterior temporal and parahippocampal cortex, followed by global efficiency, thalamic incoming strength and a leading eigenvalue of *R* (Fig. 5C).

### 3.6 A single index of hierarchical differentiation

Because the group difference is a uniform contraction, it can be summarised by one number per infant: the standard deviation of the net-asymmetry profile across the 46 regions, a template-free measure of how sharply differentiated the hierarchy is. This index was 1.62 in term-born and 0.885 in preterm infants at term-equivalent age, a ratio of 0.55 that reproduces the region-by-region slope (Hedges’ *g* = 1.51, 95% CI [1.29, 1.83]; AUC = 0.865, [0.817, 0.905]; Fig. 6A). The separation exceeds that of any individual region in Fig. 3, where the largest effect size is 1.26.

**Figure 6:**
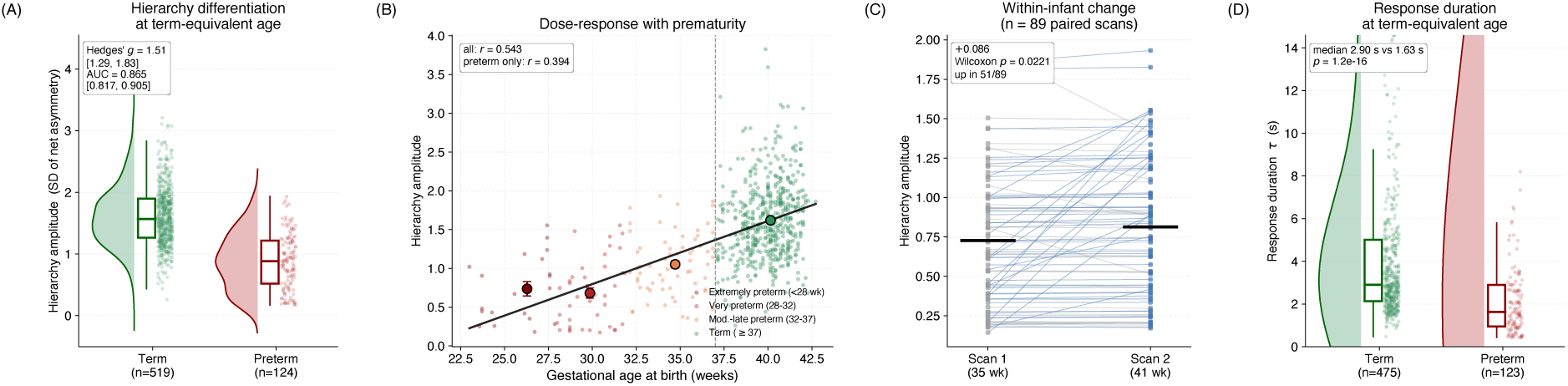
Hierarchical differentiation as a single per-infant index. **(A)** Hierarchy amplitude – the standard deviation across the 46 grey-matter regions of a subject’s net-asymmetry profile – in term-born and preterm infants scanned at term-equivalent age. The measure is template-free: it requires no reference group and is therefore not circular. Half-violins show the kernel density, boxes the interquartile range and median, dots individual infants. Effect size and discrimination are annotated with bootstrap 95% confidence intervals computed on balanced resamples (*n* = 124 per group, *B* = 2000), so neither is inflated by the 4:1 group imbalance. **(B)** The same index against gestational age at birth, coloured by degree of prematurity; the black line is the ordinary least-squares fit and the large markers give stratum means ± 1 SEM. The relationship holds within preterm infants considered alone (*r* = 0.39), and therefore does not reduce to a term/preterm dichotomy; the dashed line marks the 37-week threshold. **(C)** Within-infant change across the 89 infants imaged twice (median interval 7.3 weeks). Each line is one infant; blue lines increase. Thick bars are group means. The index does not rise significantly between sessions and, within term-born infants, it does not vary with postmenstrual age at scan (*r* = −0.05, *p* = 0.24): he behaviour expected of a maturational endpoint. **(D)** Intrinsic response duration *τ* of the slowest mode at term-equivalent age. Infants whose leading mode does not decay (Re(*λ*) ≥ 0; 45 of 643) are excluded, and the axis is truncated at the 95th percentile because the distribution has a heavy right tail; medians and inter-quartile ranges are unaffected.

The index scaled with gestational age at birth across the whole sample (*r* = 0.54) and within preterm infants considered alone (*r* = 0.39, *p* = 6 × 10^−6^), so the relationship is not a term/preterm dichotomy (Fig. 6B). By stratum it was 0.37 for extremely preterm, 0.37 for very preterm, 0.61 for moderate-to-late preterm and 1.62 for term-born infants, a graded relationship that appears to saturate below about 32 weeks. Within term-born infants the index did not vary with postmenstrual age at scan (*r* = −0.05, *p* = 0.24), yet across the 89 infants imaged twice it increased significantly between sessions (Wilcoxon *p* = 0.022; Fig. 6C).

The deficit did not attenuate across the window studied. Within preterm infants, hierarchy amplitude was unrelated to postmenstrual age at scan (*r* = −0.000, *p* = 0.998; *r* = 0.10, *p* = 0.26 after controlling for gestational age at birth), and preterm infants scanned in the later half of the window were no closer to the term-born mean than those scanned earlier (0.877 versus 0.893, against 1.620 in term-born infants). At term-equivalent age the difference is therefore persistent rather than closing.

Response duration showed the same contrast: at term-equivalent age the slowest mode persisted for a median of 2.90 s in term-born but 1.63 s in preterm infants (*p* = 1.2 × 10^−16^; Fig. 6D).

Head motion differed between groups, with term infants moving *more* (median framewise displacement 0.605 versus 0.469, *p* = 7 × 10^−8^), but motion was uncorrelated with the index (*r* = 0.012, *p* = 0.77); sex had no effect (*p* = 0.17). The index was unchanged when groups were matched one-to-one on postmenstrual age at scan (*g* = 1.51) and had a split-half reliability of *r* = 0.95 within a single scan (Appendix A).

## 4 Discussion

In recent years, resting-state functional MRI (rs-fMRI) has emerged as a powerful tool for characterizing early functional brain organization in preterm infants, providing a systems-level perspective on how prematurity alters the maturation of large-scale neural networks [33, 34]. Accumulating evidence indicates that large-scale functional networks are already detectable during the neonatal period, although their spatial coherence, integration, and temporal stability remain developmentally immature [10–13]. Within this framework, prematurity is increasingly interpreted not as a focal loss of function but as a disturbance of the developmental trajectory of large-scale network organization. This interpretation aligns with the concept of brain dysmaturation, whereby early extrauterine exposure alters the timing and coordination of neural circuit maturation rather than producing purely destructive injury [21, 35, 36]. A central finding across neonatal rs-fMRI studies is that primary sensorimotor and sensory networks emerge early in both term and preterm infants, whereas higher-order associative systems, including default-mode and frontoparietal networks, follow a more protracted developmental trajectory [10–13, 37]. In preterm populations, rs-fMRI studies have consistently reported reduced within-network connectivity, altered inter-network segregation, and less efficient global network topology relative to term-born controls [20, 34]. These alterations are increasingly interpreted within a dysmaturation framework, reflecting disrupted developmental trajectories rather than overt structural injury [21, 35, 36]. Longitudinal studies further suggest that neonatal functional connectivity carries prognostic information regarding later neurodevelopment [38]. In cohorts acquiring rs-fMRI at term-equivalent age, differences in early network organization have been associated with motor, cognitive, and behavioral outcomes during infancy and early childhood [33, 34, 39]. Altered connectivity within sensorimotor, salience, and emerging higher-order networks has been linked to later developmental performance even in infants without major structural brain injury [33, 34]. Together, these findings indicate that neonatal functional connectivity may provide complementary prognostic information beyond conventional structural MRI, although translation into routine clinical practice remains an important challenge. Methodologically, recent studies have increasingly adopted dynamic descriptions of functional connectivity that capture time-varying network organization rather than assuming stationary connectivity across the scan [15, 16, 23]. Emerging neonatal data indicate that preterm infants may exhibit a reduced repertoire and flexibility of connectivity states, potentially reflecting immature or constrained network dynamics [34]. While these approaches provide valuable insight into the temporal organization of developing networks, they remain sensitive to motion, scan duration, and analytic choices, highlighting the need for methodological standardization before widespread clinical application. Against this background, the present study introduces a response-based framework derived from statistical physics to quantify directed functional responsiveness in neonatal brain networks. Unlike conventional functional connectivity, which captures undirected correlations between brain regions, this approach estimates how activity in one region would dynamically respond to perturbations in another, thereby providing a temporally informed measure of directed influence. By integrating directional response properties with conventional connectivity measures, this framework enables a more comprehensive characterization of the developing functional connectome. Using this approach, we observed that the dynamic response metric exhibited substantially larger relative changes across gestational age bins than conventional functional connectivity. While FC increased gradually with advancing age, the response-based metric displayed a steeper developmental trajectory, particularly during earlier gestational windows (Fig. 1). This finding suggests that temporally informed descriptors of BOLD dynamics may provide enhanced sensitivity to maturational processes occurring during the late preterm and early term-equivalent period. Previous neonatal rs-fMRI studies have shown that conventional FC primarily reflects the progressive strengthening and consolidation of large-scale network architecture [10–13]. However, increasing evidence indicates that early brain development involves rapid changes not only in connectivity strength but also in the temporal organization and flexibility of network interactions [15, 16, 23]. Dynamic connectivity analyses have revealed developmental effects that are partially dissociable from static FC, including age-related increases in network-state variability and repertoire complexity [34]. Our finding that response-based dynamics are particularly sensitive to gestational age differences is therefore consistent with a broader framework in which early brain development is characterized by the progressive reorganization and maturation of transient functional network dynamics.[34, 40] One plausible interpretation is that dynamic response metrics capture processes such as the stabilization of oscillatory regimes, refinement of neurovascular coupling, or increasing efficiency of network switching, which may precede detectable changes in average connectivity strength. This interpretation is supported by recent work demonstrating that the variability and temporal structure of neonatal BOLD signals evolve rapidly during the preterm period and may provide early markers of brain dysmaturation [9, 34]. Beyond global maturation effects, our analysis revealed a clear developmental dissociation between outgoing (driver strength) and incoming (listener strength) influences across postmenstrual age (Fig. 2). Outgoing influence increased progressively, particularly within parietal regions, whereas incoming sensitivity remained relatively stable or declined in several areas. These changes resulted in increasingly negative net information flow in selected hubs, suggesting the progressive emergence of hierarchical organization within the developing functional connectome. This pattern is consistent with contemporary models of brain network development proposing a transition from relatively symmetric and locally organized interactions toward more hierarchical and hub-dominated architectures [15, 16, 41]. Connectome studies have shown that late gestation is characterized by rapid increases in network integration and the consolidation of hub regions within posterior associative cortices [20, 35, 41]. Effective connectivity analyses further indicate that maturational changes are often more pronounced in measures capturing directed influence or information flow than in undirected functional connectivity alone [23]. The prominent increase in outgoing influence observed in parietal regions aligns with prior evidence for a posterior-to-anterior gradient of functional maturation. Several neonatal imaging studies have demonstrated earlier maturation of posterior cortical systems, including parietal and sensorimotor regions, followed by more protracted development of frontal control networks [10–13, 35]. Within this framework, the parietal cortex may function as an early integrative scaffold of the developing connectome, progressively increasing its broadcasting capacity as long-range cortico-cortical and thalamocortical pathways mature. Conversely, the relative stability or reduction in incoming sensitivity observed in some regions may reflect increasing functional selectivity and network refinement. Early neonatal networks are characterized by diffuse and redundant coupling, which gradually gives way to more efficient and specialized communication patterns as development progresses [15, 16]. Our analyses also revealed structured hemispheric asymmetries in dynamic response metrics, particularly within temporal and cingulate regions, as well as differential lateralization patterns between term and preterm infants (Fig. 4). These findings indicate that hemispheric specialization is already emerging during the late preterm and early term-equivalent period. Previous neonatal imaging studies have similarly documented early hemispheric biases within perisylvian and temporal networks, suggesting that proto-language circuits begin to organize before full-term maturation [18–20]. Importantly, the dynamic response framework extends these observations by demonstrating that hemispheric asymmetry is not limited to undirected connectivity strength but also affects the directionality of information flow. Differences in driver versus listener roles between hemispheres suggest that early functional specialization may involve asymmetric maturation of broadcasting and integrative functions across cortical systems. Strikingly, in seven regions the two directions are lateralised in *opposite* senses, so that a region may be recruited into one hemispheric system for the influence it receives and into the other for the influence it exerts. The posterior superior temporal gyrus, a core node of the emerging language scaffold [42], listens right-dominantly while driving left-dominantly, which suggests that the left-lateralisation of perisylvian language circuitry may be established first in the outgoing direction. Prematurity was associated with measurable alterations in these lateralization patterns, particularly within cingulate and temporal territories. Disruption of typical hemispheric specialization has previously been reported in preterm populations across both structural and functional modalities and has been linked to altered white matter development and later language and cognitive vulnerabilities [21, 24]. This finding may be particularly relevant in the context of the well-established vulnerability of language development following preterm birth. Meta-analytic evidence indicates that preterm-born children show poorer performance than term-born peers across both simple and complex language functions, with differences in complex language abilities becoming increasingly apparent across childhood [43]. More recent evidence indicates that differences in both receptive and expressive language are already detectable within the first 18 months of life, with lower gestational age being associated with greater differences in receptive language performance [44]. Given the central role of the superior temporal cortex in the emerging language network, altered directional lateralization in this region may therefore represent an early systems-level signature of atypical specialization within circuits supporting later language development.Given the central role of the superior temporal cortex in early language network scaffolding, altered lateralization patterns in preterm infants may therefore reflect early disturbances in hemispheric specialization processes relevant to later neurodevelopment. At the network level, our findings further indicate that prematurity is associated with an attenuation, rather than a rearrangement, of hierarchical information flow. In term-born neonates, associative cortical regions exhibited driver-dominant profiles while subcortical and limbic structures were consistently listener-dominant, consistent with early emergence of distributed cortical–subcortical control architectures. In contrast, preterm infants showed attenuation of driver properties in parietal, frontal, temporal, and occipital regions, accompanied by an equally proportionate weakening of the listener role of thalamus, amygdala and basal ganglia. The regularity of this effect is its most informative feature (Fig. 3 and 6). Every region retained its position in the hierarchy, and a single multiplicative factor of about one half recovers 98% of the preterm profile from the term profile. Prematurity therefore does not appear to redistribute computational roles between cortical and subcortical territories, nor to invert the cortical–subcortical axis; it leaves the blueprint of the hierarchy intact while failing to deepen it. This remarkably preserved yet attenuated hierarchical organization suggests that preterm birth leaves a measurable imprint on the developing functional connectome that remains detectable at term-equivalent age. This imprint does not appear to take the form of a fundamentally different network architecture: the overall organization is preserved, but its functional differentiation is attenuated. This interpretation is consistent with previous resting-state fMRI studies showing that, despite ongoing maturation, very preterm infants continue to exhibit reduced network integration, segregation and connectivity strength compared with term-born neonates at comparable postmenstrual ages[45–47]. Thus, reaching term-equivalent age does not necessarily imply functional equivalence to being born at term. Rather, premature exposure to the extrauterine environment appears to leave a persistent and measurable signature on the developing brain.

Read alongside the developmental trajectories, preterm infants at term-equivalent age resemble an earlier point on the same maturational axis, but one they are not visibly leaving: within preterm infants scanned between 37 and 46 weeks, hierarchy amplitude is unrelated to postmenstrual age at scan, and those scanned later are no closer to the term-born mean. The sign is persistent rather than transient. A preterm infant imaged at term-equivalent age is therefore not equivalent to a term-born infant of the same postmenstrual age, even though term-equivalent imaging is routinely treated as the point at which the two become comparable. This is the disturbance of developmental timing originally proposed for preterm brain injury [36], here reduced to a single structural quantity,one that, unlike machine-learning estimates of brain maturity [21, 37], is a direct property of the directed hierarchy rather than the output of a trained predictor

The shortening of intrinsic response duration in preterm infants points in the same direction. A brain whose directed hierarchy is shallower also relaxes faster after perturbation, so reduced responsiveness and shortened persistence are not independent deficits but two readings of the same under-differentiated architecture. Longer intrinsic timescales in mature cortex are generally taken to reflect the capacity to integrate information over extended windows [48]; on that view, the halving of response duration after preterm birth would represent a reduced temporal integration window at an age when sensory and associative systems are beginning to be recruited by experience. Finally, our predictive modeling demonstrated that directed functional network features at scan age contain substantial information about birth timing, achieving moderate predictive performance (*R*^2^ = 0.50 from network features alone, rising to *R*^2^ = 0.60 once postmenstrual age at scan is included). The most informative predictors were the incoming strength of the parietal lobe bilaterally and the outgoing strength of anterior temporal and parahippocampal cortex, followed by global efficiency, thalamic incoming strength and a leading eigenvalue of *R* (Fig. 5 These regions are known to play central roles in early connectome integration and to be particularly vulnerable to altered developmental trajectories in preterm populations [20, 35, 49]; notably, both node-level directionality and a global timescale descriptor contribute, indicating that prematurity is encoded at more than one level of the response architecture. From a systems neuroscience perspective, these findings support the concept of a distributed “network chronotype”, whereby early functional architecture retains measurable signatures of neurodevelopmental timing. Prior studies have demonstrated that neonatal functional connectivity patterns can predict developmental age and characterize early maturation trajectories [21, 37]. Our results extend this framework by showing that directed, temporally informed metrics capture birth-related variance beyond what is typically observed using static connectivity measures alone. Together, these findings indicate that prematurity leaves a measurable imprint on the directional architecture of the neonatal connectome. By capturing hierarchical information flow, hemispheric specialization, and dynamic responsiveness, response-based metrics provide a sensitive systems-level marker of early brain maturation and reveal how prematurity reshapes the developing functional connectome.

## 5 Conclusions

By integrating static and dynamic measures of functional organization, this study provides a comprehensive characterization of neonatal brain maturation from preterm to term age. Our findings demonstrate that functional development is reflected not only in progressive changes of connectivity strength, but also in evolving patterns of responsiveness, directed information flow, and hemispheric specialization. While conventional functional connectivity shows gradual age-related modulation, dynamic response metrics exhibit faster and more pronounced developmental sensitivity, highlighting their potential as markers of neurodevelopmental maturity (Fig. 1). Developmental trajectories further reveal a progressive reorganization of driver and listener roles across brain regions, accompanied by increasing net information flow and stabilization of local hub architecture over time (Fig. 2). Importantly, term- and preterm-born neonates display distinct patterns of functional lateralization, indicating early divergence in hemispheric specialization (Fig. 4). At term-equivalent age, preterm infants retain an altered balance between incoming and outgoing information flow, defining a network-level dysmaturity signature that persists despite comparable postmenstrual age (Fig. 3). Condensed into a single per-infant index, this signature separates the groups more sharply than any individual region, scales with gestational age at birth, and is accompanied by a halving of intrinsic response duration (Fig. 6). Taken together, these results suggest that prematurity leaves a measurable imprint on the hierarchical and dynamical organization of the neonatal connectome, and that response-based metrics provide a sensitive framework for detecting early deviations in functional brain development. What that imprint appears to be is not a rewiring of the developing connectome but a failure to complete its differentiation. This persistent and quantifiable signature may therefore represent a potential biomarker of functional brain maturation, providing a sensitive means of identifying early deviations from typical neurodevelopmental trajectories.

## Appendix: supplementary information

## A Robustness of the hierarchy-differentiation index

### Definition and the circularity it avoids

Two related indices can be built from a subject’s net-asymmetry profile *v* ∈ R^46^. The first projects it onto the term-group mean profile *v̄*_term_, HDI = *v, v̄*_term_ */ v̄*_term_*, v̄*_term_, which is convenient because HDI = 1 denotes a fully term-like hierarchy, but which is mildly circular: the term group defines the reference against which it is then measured, so its mean is exactly 1 by construction. A leave-one-out reference shows the circularity is numerically negligible at this sample size (term mean 0.9998, Hedges’ *g* unchanged at 1.53). We nonetheless report as primary the *template-free* alternative, the standard deviation of *v* across regions, which requires no reference group at all. The two agree almost perfectly (*r* = 0.992), and all conclusions are identical under either.

### Class imbalance

The term and preterm groups differ in size by a factor of four. Hedges’ *g* already carries the unequal-*n* correction and the area under the ROC curve is prevalence-independent, so neither statistic is biased by the imbalance; we verified this directly.

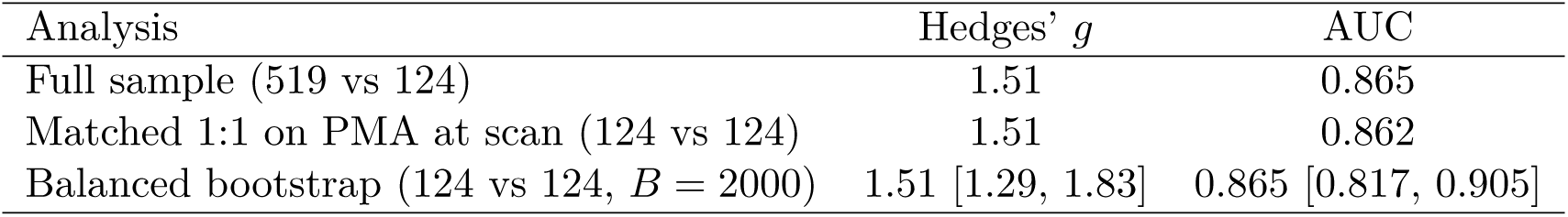

Postmenstrual age at scan does not differ between the groups within the 37–46 week window (median 41.29 vs 41.00 weeks, *p* = 0.28), so matching changes little. The matched estimate is a point estimate rather than an interval: greedy nearest-neighbour matching selects nearly the same term infants on every draw. We emphasise effect sizes over *p*-values throughout, because with *n* = 643 the latter (of order 10^−37^) reflect sample size rather than effect magnitude.

### Motion and sex

Head motion differs between groups, with term infants moving *more* (median framewise displacement 0.605 vs 0.469, *p* = 7 × 10^−8^). Motion is nevertheless uncorrelated with the hierarchy index (*r* = 0.012, *p* = 0.77), so the group effect cannot be motion-driven and the direction of the motion difference is opposite to the one that would be needed to produce it. Sex has no effect on the index (*p* = 0.17).

### Within-scan reliability

Splitting each acquisition into its first and second halves and recomputing the index independently gives *r* = 0.949 between halves (*n* = 150; Spearman–Brown corrected 0.974), so the measure is stable within a single 15-minute scan.

## B Sensitivity to the regularisation constant

The Tikhonov constant *λ* in Eq. (3) was set to 10^−5^. Recomputing the index over four orders of magnitude leaves it essentially unchanged; only at *λ* = 10^−2^, where the regulariser becomes comparable to the diagonal of *C*^^^, does appreciable shrinkage appear (subsample of *n* = 200 at term-equivalent age).

| $\lambda$ | Term | Preterm | Ratio | $r$ with $\lambda = 10^{-5}$ |
| --- | --- | --- | --- | --- |
| $10^{-6}$ | 1.617 | 0.863 | 0.534 | 1.0000 |
| $10^{-5}$ | 1.616 | 0.863 | 0.534 | — |
| $10^{-4}$ | 1.611 | 0.862 | 0.535 | 1.0000 |
| $10^{-3}$ | 1.563 | 0.851 | 0.544 | 0.9992 |
| $10^{-2}$ | 1.258 | 0.761 | 0.604 | 0.9737 |

## C Validity of the first-order Jacobian

The estimator *J* = (*R − I*)*/*Δ*t* is the first-order expansion of *R*(Δ*t*) = exp(*J* Δ*t*) and is accurate only for modes whose eigenvalue lies near unity. Across subjects the eigenvalues of *R* are strongly bimodal: the leading eigenvalue has median 0.869 (IQR 0.813–0.931), and a median of exactly five modes exceed 0.5, while the remaining forty-one cluster near zero (median *λ* = 0.127). The five leading modes reported in Fig. 1C are therefore precisely the modes for which the expansion is justified; the fast bulk of the spectrum is not interpreted.

Comparing the first-order timescale *τ* = Δ*t/*(1 − *λ*) against the exact matrix-logarithm solution *τ* = − Δ*t/* ln *λ* for the leading mode gives medians of 2.86 s and 2.66 s respectively: the first-order form overestimates *τ* by a median of 7.6% (IQR 4.4–11.9%). The two are rank-identical (*r* = 1.0000), so every group comparison, correlation and developmental trajectory reported here is unchanged in significance or direction; only the absolute timescale carries this modest upward bias.

In 8% of subjects the leading eigenvalue equals or exceeds unity, so the mode does not decay and *τ* diverges. Such subjects are assigned *τ* = +, which preserves their rank as maximally persistent, and all summaries of *τ* use the median rather than the mean for this reason; the mean is not merely noisy but can be negative (see Fig. 1C).

## D Global response magnitude versus hierarchical differentiation

Preterm infants show both a lower overall magnitude of directed response and a flatter hierarchy, and the two are strongly collinear (*r* = 0.96 between total directed strength and hierarchy amplitude). Distinguishing them matters, because a uniform change of gain would produce a reduced amplitude without any change in hierarchical organisation.

Total absolute directed strength is 147.1 in term-born and 115.8 in preterm infants (ratio 0.787, *g* = 1.50), against 1.620 and 0.885 for hierarchy amplitude (ratio 0.546, *g* = 1.51). The amplitude therefore falls further than the total. Dividing each infant’s profile by their own total strength yields a scale-free measure that is invariant to any global gain: the group difference persists and increases (ratio 0.673, *g* = 1.65, *p* = 6 × 10^−33^). The flattening is thus disproportionate to the overall reduction, by approximately a third.

We note one analysis that appears to conflict with this and should not be used. Residualising amplitude linearly on total strength across the pooled sample reduces the group difference to *g* = 0.20 (*p* = 0.07). This is an artefact of regressing out a covariate that itself differs strongly between the groups and is 96% correlated with the measure of interest: the regression absorbs the very effect under test. The ratio normalisation above is the appropriate control, since it is invariant to gain by construction rather than by estimation.

## E Region selection

The dHCP neonatal parcellation contains 87 structures. All analyses use the 46 cortical and subcortical grey-matter regions that remain after excluding white matter, CSF, ventricles, corpus callosum, brainstem, cerebellum and background, matched case-insensitively on structure name. The retained set is exactly bilateral (23 left, 23 right), which is what makes the 23 homologous pairs of Fig. 4 well defined. The same 46 regions are used for every figure, so region-level results are directly comparable across analyses.

## F Reproducibility

All analyses were run in Python 3.11 (NumPy, SciPy, scikit-learn, statsmodels, nibabel). A single notebook performs the complete analysis and writes every figure in the paper, reading the region set, the time-series loader and the response estimator from one shared configuration block so that region indexing, header handling and the value of *λ* cannot differ between analyses.

